# An extracellular matrix coating similar to articular cartilage inhibits the differentiation of myofibroblasts and limits the development of implant-induced fibrosis

**DOI:** 10.1101/2025.09.10.675333

**Authors:** Melis Ozkan, Michael JV White, Ani Solanki, Darrell Pilling, Richard H Gomer, Jeffrey A Hubbell

## Abstract

Fibrosis is involved in 45% of deaths in the United States, with no treatment options to reverse progression of the disease. Implantable devices (such as joint replacements, chips, stents, artificial organs, biosensors, catheters, heart valves, scaffolds for tissue engineering, etc.) can trigger a foreign body response, in which fibrotic tissue covers the implant and impedes its function. Myofibroblast are a key cellular component of scar tissue. To explore the relationship between extracellular matrix-based coatings and fibrosis, we coated tissue-culture surfaces using a library of extracellular matrix (ECM) proteins and then performed an in-vitro screen for myofibroblast differentiation on the coated surfaces as an indicator of fibrotic potential. The protein and proteoglycan components of cartilage (collagen II, biglycan, decorin, and chondroitin sulfate) were individually anti-fibrotic. Further, mixtures of collagen II, biglycan, decorin, and chondroitin sulfate inhibited myofibroblast differentiation to a greater degree than collagen II, biglycan, decorin, or chondroitin sulfate as individual coatings. Next, we performed an *in-vivo* model of a foreign-body response. Implanting an uncoated implant subcutaneously into mice resulted in a thicker layer of fibrotic scar tissue than implants coated with a cartilage-like ECM mixture. Our results indicate that the ECM microenvironment is key to the initiation, progression, and maintenance of fibrosis.

## Introduction

Scar tissue deposition is the primary cause of both the foreign-body response and fibrosing diseases [1]. The foreign body response involves the innate immune system and nearby recruited cells encapsulating the foreign object in a fibrous sheath [2], similar to a desmoplastic response of fibrotic tissue surrounding tumors. While desmoplastic responses can encapsulate structures of multiple origins including several parasites [3-6] and multiple types of tumors [7-12], the foreign body response is a result of implantable devices [2] which are almost always considerably stiffer than biological tissues. The resulting fibrotic sheath responds to changing conditions in its microenvironment, including mechanical forces, enzymatic digestion, scar tissue-hydration, inflammation, and ECM-crosslinking [13, 14]. The global expense for implantable devices is expected to reach $155B by 2026 [15]. 30-50% of pacemakers, and 30% of mammary implants, will require replacement over their lifespan due to implant fibrosis caused by the foreign-body response [16]. A conservative estimate of a 10% replacement rate for implants due to the foreign-body response indicates the cost to the US healthcare system could be $10B annually [16].

Fibrosing diseases such as pulmonary fibrosis, congestive heart failure, liver cirrhosis, and end-stage kidney disease occur when scar tissue develops in or on an otherwise healthy organ. Fibrosis is involved in up to 45% of deaths in the United States [17, 18]. The etiology of fibrosis is poorly understood [17], and current treatment options are limited. The only three treatments that are approved for fibrosis slow, but do not reverse, the progression of the disease [18-21].

Myofibroblasts are key to scar tissue development. Myofibroblasts are defined by their functions (actively depositing extracellular matrix (ECM), secreting pro-fibrotic signals, and by physically stiffening tissues through tension force generated by their cytoskeleton [17, 22, 23]) and by their spindle-shaped morphology, rather than by cell lineage [24]. Myofibroblasts can arise from several progenitors including hepatocytes, fibroblasts, epithelial cells, and monocytes [17, 24, 25]. Myofibroblast contractility is key to the progression of fibrosing diseases, and myofibroblasts can be activated by tissue stiffness [26]. Myofibroblasts are key to the development of implant fibrosis in the foreign-body response [1].

Mechanosensing is a dynamic process integrating multiple signals between cell surface integrins, intracellular machinery, and secreted signals. Mechanosensing is key to the development and maintenance of the myofibroblast phenotype [22, 23]. Mechanosensing by monocyte-derived cells and myofibroblasts is key to the development of the foreign-body response [27].

The intact ECM of healthy, undamaged tissues “stress-shields” cells from tension forces, with the intact ECM protecting cells from tension forces. This is why myofibroblasts are formed or recruited in response to injury which damages tissue, disrupting the intact ECM architecture that had previously “stress-shielded” cells [25]. By reducing the stiffness of the environment, normal healthy tissues reduce myofibroblast activation [28]. Among myofibroblast precursors, monocytes are unique in that they can be recruited from the bloodstream to different tissues in the body [29]. These monocyte-derived myofibroblasts may provide a third of myofibroblasts in liver fibrosis [30], kidney [31], lung [32, 33], and skin fibroses [34].

Cartilage is a resilient, flexible tissue that performs multiple structural roles in the body. Cartilage is composed of specialized cells called chondrocytes, surrounded by a network of ECM largely composed of collagens and proteoglycans [35]. Cartilage is heterogenous and varies in its ECM composition, in the ratio of proteoglycans and proteins that make up the cartilage, and in the hydration state of the matrix. These variances cause cartilage to have different properties. For instance, an osteoarthritic cartilage decreases its stiffness while increasing in hydration [36, 37].

Articular cartilage is found in joints, primarily serving to absorb the compressive and shear forces of load bearing joints resulting from movement [38]. Articular cartilage is composed of four stress-diffusing layers (superficial, intermediate, deep, and calcified) with the protein composition changing between layers [35, 38]. Articular cartilage interacts with the synovial fluid in the middle of the joint and is primarily composed of several different kinds of ECM including collagen II, chondroitin sulfate, biglycan, and decorin [35]. Within cartilage, different layers and mixtures of ECM form a complex, dynamic environment that varies greatly in ability to withstand compressive forces and strain [35].

Collagens are a family of fibrillar proteins composed of α-chain trimers, the most common type of protein found in the human body, and are highly evolutionarily conserved [39]. Collagens are defined by their quaternary trimeric structures, formed by the interaction of three α-chains that differ among collagen subtypes [39]. Collagen II is the type of collagen found in cartilage [35, 39] and is composed of three α1(ll) chains [40].

Biglycan is a small leucine-rich proteoglycan (SLRP) that is expressed ubiquitously and serves several structural and signaling roles in the body [41]. Biglycan interacts with transforming growth factor beta (TGFβ), tumor necrosis factor alpha (TNFα), collagens I-IV, Toll-like receptors 2 and 4 (TLR2, 4), and low-density lipoprotein receptor-related protein 6 [41]. Biglycan is thus an innate immune system signaling molecule, in addition to its role in the ECM [42], and can mediate between inflammatory and anti-inflammatory responses depending on context [43].

Decorin is a SLRP closely related to biglycan and lumican [44] that canonically binds to and acts in the organization of collagens [45]. Like biglycan, decorin has many roles including interactions with collagens and other components of the ECM, signaling receptors, and ligands [46]. Interestingly, decorin’s (and biglycan’s) ability to bind collagens can enhance the ability of a collagen-containing matrix such as cartilage to bear stresses [44].

Chondroitin sulfate (CS) is a glycosaminoglycan (GAG) usually found attached to matrix proteins as part of a proteoglycan [47] or a protein that is heavily glycosylated [47]. Chondroitin sulfate is an important structural component of cartilage [35].

Here we show that monocytes cultured on collagen II, biglycan, decorin, or chondroitin sulfate resist differentiation to a myofibroblastic phenotype. Further, collagen II, biglycan, decorin, or chondroitin sulfate form a more anti-fibrotic culture surface when mixed. These findings are of interest not only for the possibility of creating anti-fibrotic coatings for implantable devices, but may explain why cartilage, one of the stiffest hydrated surfaces in the human body, tends to be free of fibrotic tissue.

## Results

We began our studies with the observation that monocyte differentiation into myofibroblasts was dependent on adhesion to a stiff surface [48, 49]. For this study, we expanded into an exploration of whether monocyte-to-myofibroblast differentiation was also governed by the protein composition of that stiff surface.

To determine which, if any, ECM proteins affected monocyte-to-myofibroblast differentiation, we performed an in-vitro screen. We placed freshly purified primary monocytes from human blood donors onto tissue-culture treated plastic coated with one ECM protein. We allowed the monocytes to differentiate into myofibroblasts over the course of 5 days, and counted the number of spindle-shaped, morphological macrophages, which indicate a myofibroblast-like state (Fig. 1). We normalized the myofibroblast counts for each donor to the monocyte-myofibroblast count on uncoated tissue-culture treated plastic to account for a previously observed wide range of donor monocyte variability.

**Figure 1:**
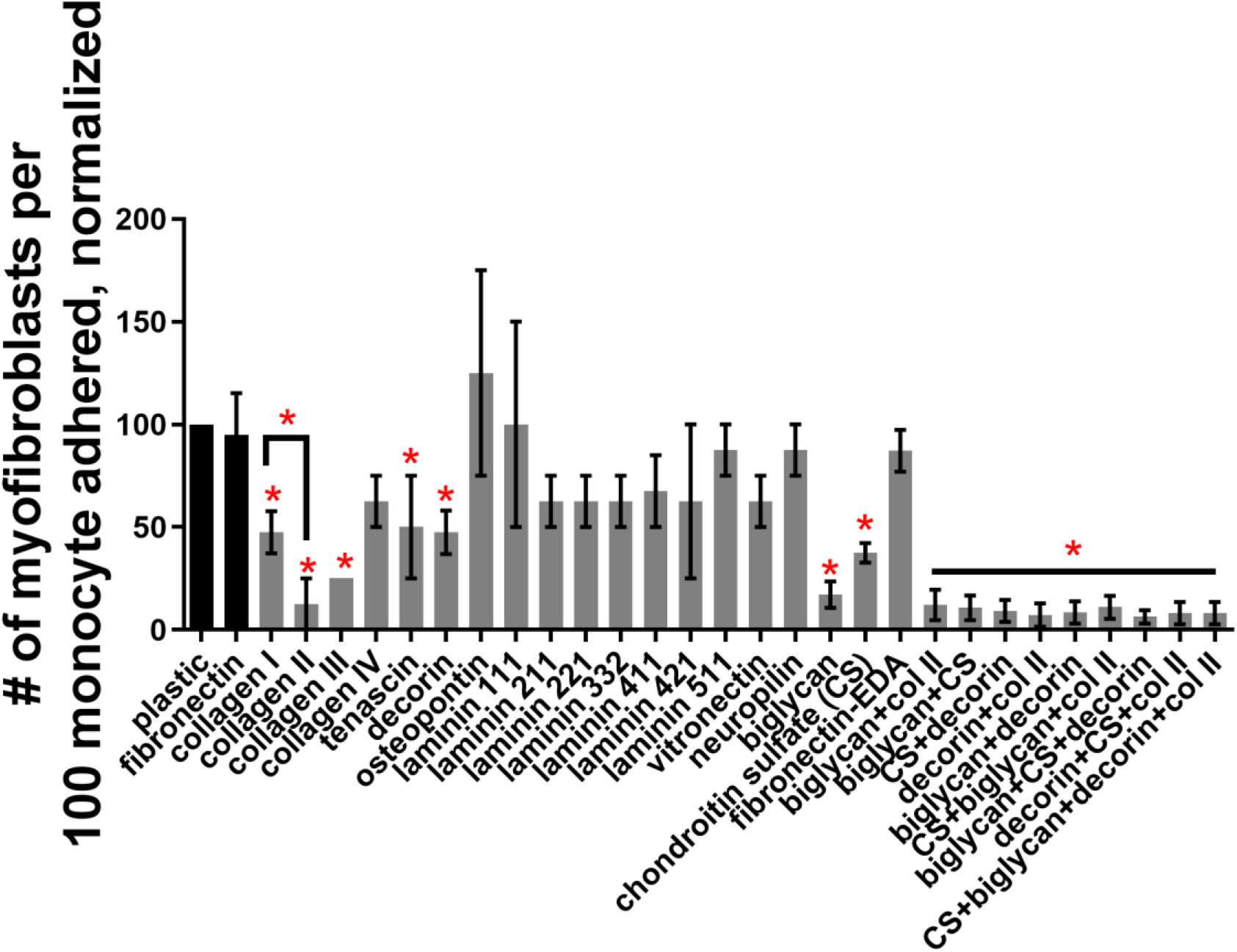
Several ECM coatings individually inhibit the differentiation of monocytes into myofibroblasts, and the combination of ECM components of cartilage also inhibit monocyte-to-myofibroblast differentiation. Freshly isolated human monocytes were cultured on 24 well tissue culture plates coated with one or more ECM proteins. Myofibroblast differentiation was assessed by morphology, counting the number of spindle-shaped myofibroblasts as a percentage of plated monocytes. Fibronectin-EDA is the fibronectin splice variant containing the extra-domain A. Chondroitin sulfate is denoted as CS. Values are mean ± SEM, n ranging from 2 to 9. * P < 0.05 compared to fibronectin control, 1-way ANOVA with Dunnett’s post test. Figure 1 is significant by 1-way ANOVA with a post-test for trend between datasets read in a left-to-right direction.

Several ECM components inhibited the formation of monocyte-derived myofibroblasts, including collagens I, II, III, tenascin, decorin, biglycan, and chondroitin sulfate (Fig. 1). Collagen II inhibited monocyte-myofibroblast differentiation more than collagen I. Collagen II, decorin, biglycan and chondroitin sulfate are the primary components of cartilage. A surface coating mixture of collagen II, decorin, biglycan and chondroitin sulfate, where the relative proportions of the four components approximate the proportions found in healthy cartilage [35], was most inhibitory of monocyte-myofibroblast differentiation (Fig. 1). Interestingly, there is a significant trend for lower monocyte-myofibroblast differentiation in Fig. 1, as measured by a 1-way ANOVA with a post-test for trend, which tests for a linear trend among datasets read from a left-to-right direction. Put another way, fewer myofibroblasts differentiate on average on surfaces coated with collagen II, decorin, biglycan or chondroitin sulfate when compared with the myofibroblast differentiation on uncoated plastic, and fewer still myofibroblasts differentiate on the mixtures of collagen II, decorin, biglycan or chondroitin sulfate, and this trend is significant. This conforms with our observation that increasingly cartilage-like surface coatings reduce the myofibroblasts phenotype of cultured cells.

Myofibroblasts also produce more alpha smooth muscle actin (αSMA) and collagen I than do monocytes [22, 50]. To confirm that we were truly seeing a reduction in myofibroblast differentiation and not simply a reduction in spindle-shaped morphology, we repeated the coating study from Fig. 1 with a focus on the protein components of cartilage. We cultured primary human monocytes on (1) tissue-culture treated plastic or a fibronectin coating (2) tissue-culture treated plastic coated with one of collagen II, decorin, biglycan, chondroitin sulfateor (3) tissue-culture treated plastic coated with mixtures of collagen II, decorin, biglycan, and chondroitin sulfate. We then assessed the amount of myofibroblast differentiation from human monocytes as measured by flow cytometry for double-positivity for αSMA and collagen I (Fig. 2, Supplemental Fig. 1A). Culturing primary human monocytes on tissue-culture treated plastic resulted in higher numbers of αSMA and collagen I double-positive cells when compared to culturing primary human monocytes on one of collagen II, decorin, biglycan, chondroitin sulfate. Culturing primary human monocytes on mixtures of collagen II, decorin, biglycan, and chondroitin sulfate resulted in the lowest number of αSMA and collagen I double-positive cells. As in Fig. 1, there is a statistically significant trend for lower numbers of αSMA and collagen I double-positive cells when examining Fig. 2A from left to right, as measured by a 1-way ANOVA with a post-test for trend, which tests for a linear trend among datasets. This conforms with our hypothesis that increasingly cartilage-like surface coatings reduce the myofibroblasts phenotype of cultured cells.

**Figure 2:**
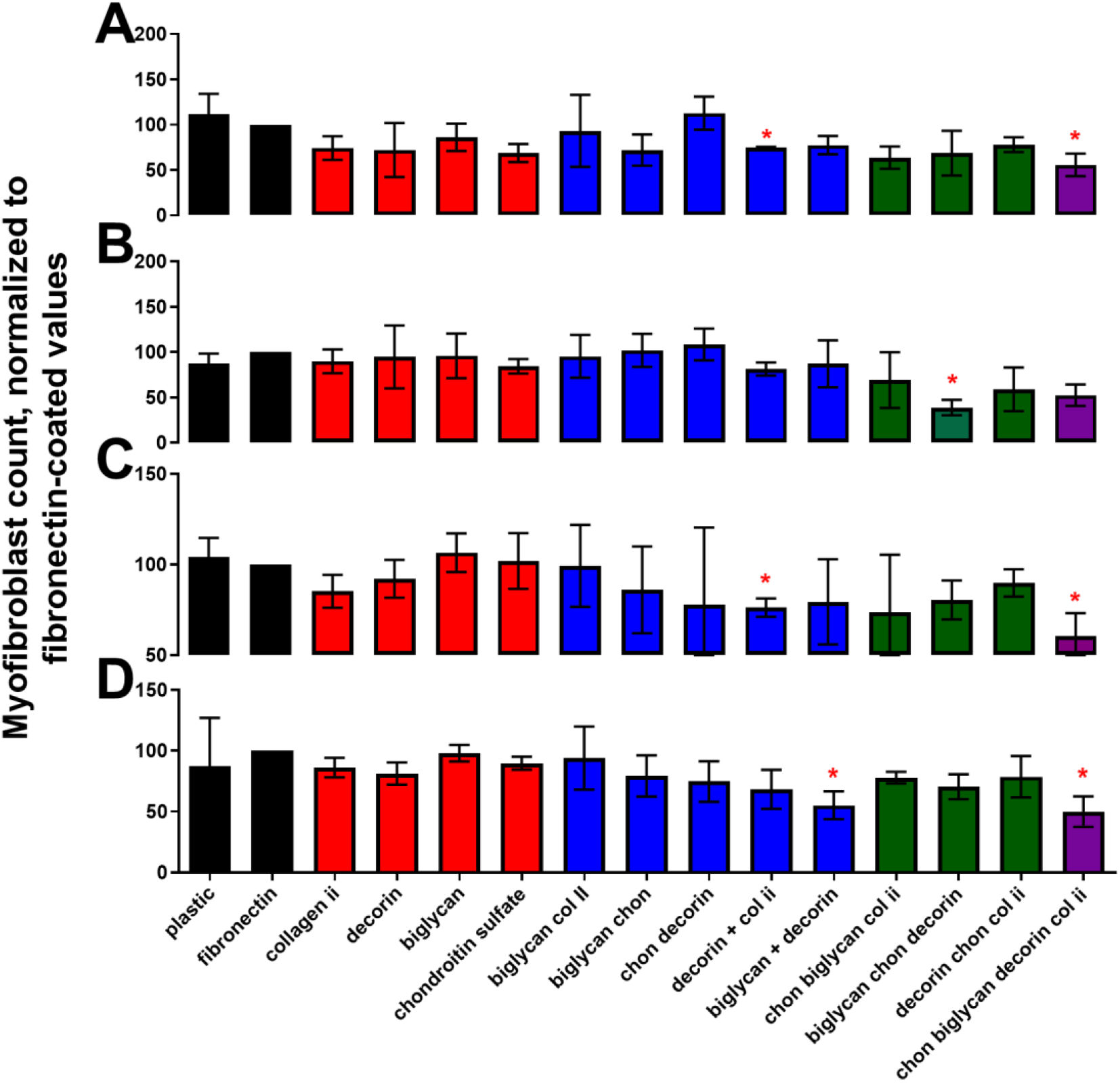
Several ECM coatings individually inhibit the differentiation of monocytes into myofibroblasts, and the combination of ECM components of cartilage also inhibit monocyte-to-myofibroblast differentiation. Freshly isolated (A) human monocytes, (B) mouse monocytes, (C) human fetal fibroblasts and (D) mouse fetal fibroblasts were cultured on 24 well tissue culture plates coated with one ECM protein (red), two ECM proteins (blue), 3 ECM proteins (green), or four ECM proteins (purple) comprising the main protein components of cartilage. Cells were allowed to become myofibroblasts over 5 days. Mouse monocyte-to-myofibroblast differentiation was induced by the addition of M-CSF and IL-13 to culture media, and fibroblast-to-myofibroblast differentiation was induced by TGFβ. Cells were removed from the plate by use of trypsin and a policeman, and assessed via flow cytometry for the percentage of the population that were αSMA and collagen I double positive. Values are mean ± SEM, n ranging from 3 to 5. * P < 0.05 compared to fibronectin coated control, 1-way ANOVA with Dunnett’s test for multiple comparisons. Each A-D panel is significant by 1-way ANOVA with a post-test for trend between datasets read in a left-to-right direction.

These findings confirmed that the mixed coating of collagen II, decorin, biglycan, and chondroitin sulfate were anti-fibrotic as coatings, based on both cellular morphology (a reduction in the spindle-shaped myofibroblast phenotype, Fig. 1) and the reduction in the amounts of proteins associated with myofibroblasts (αSMA and collagen I, Fig. 2A).

To determine if this anti-fibrotic effect extended to cells other than primary human monocytes, we also cultured freshly purified primary mouse monocytes and immortal human (MRC-5) and mouse (NIH 3T3) fibroblast cell lines on collagen II, decorin, biglycan, chondroitin sulfate, or mixtures of these proteins (Fig. 2B-D, Supplemental Fig. 1B-D). Myofibroblast differentiation was inhibited in mouse monocytes, human fibroblasts, and mouse fibroblasts when cultured with a mixture of collagen II, decorin, biglycan, and chondroitin sulfate. As with Fig. 1 and Fig. 2A, culturing cells on collagen II, decorin, biglycan, chondroitin sulfate (and mixtures of these proteins) results in a statistically significant trend of lower amounts of αSMA and collagen I in cells, as measured by a 1-way ANOVA with a post-test for trend (Fig. 2B-D). Put another way, there is a statistically significant trend towards a lower myofibroblast phenotype when reading each of Fig. 2B, 2C, and 2D from left to right.

Myofibroblasts are defined not only by their morphology (Fig. 1) and protein composition (Fig. 2, Supplemental Fig. 1), but also by their secretome. To determine if monocyte-myofibroblast differentiation is inhibited, we performed a secretome ELISA on conditioned media from human monocytes, mouse monocytes, human fibroblasts, and mouse fibroblasts cultured on collagen II, decorin, biglycan, chondroitin sulfate, or mixtures of these proteins, using a LegendPlex inflammatory signals kit (Supplemental Fig. 2).

No ECM surface coating significantly decreased the amount of IL-10 from mouse monocytes (Supplemental Fig. 2A). There was a statistically significant trend towards a decrease in concentration for CCL22, IL12-p70, IL-6, and IL-1β (Supplemental Fig. 2C, D, E, I) from mouse monocytes cultured on mixtures of ECM proteins approximating cartilage (collagen II, decorin, biglycan, chondroitin sulfate), as measured by a 1-way ANOVA with a post-test for trend. There was no significant trend for IL-10 (Supplemental Fig. 2A)

No ECM surface coating significantly decreased the amount of IL-10 from mouse fibroblasts, nor was there a significant trend for a change in IL-10 concentration resulting from culture on any individual ECM protein or proteoglycan, or mixture (Supplemental Fig. 3A). There was a significant trend for decreasing concentrations of CXCL1, IL-12p70, and IL-6, and CCL17 in mouse fibroblasts cultured on mixtures of collagen II, decorin, biglycan, chondroitin sulfate assessed by a 1-way ANOVA with a post-test for trend. The data from human monocytes and fibroblasts was below the detection limit for the LegendPlex ELISA.

To determine if a mixture of collagen II, decorin, biglycan, and chondroitin sulfate approximating cartilage would be anti-fibrotic when implanted *in vivo*, we coated two separate implants and placed them under the skin of C57Bl6 mice for 6 weeks (Fig. 3). Cartilage is comprised of four zones. The superficial zone that synovial fluid interacts with is composed of approximately 50% collagen II, 35% chondroitin sulfate, and 7.5% biglycan and decorin [35]. We coated our implants in ratios of protein identical to these superficial zone values.

**Figure 3:**
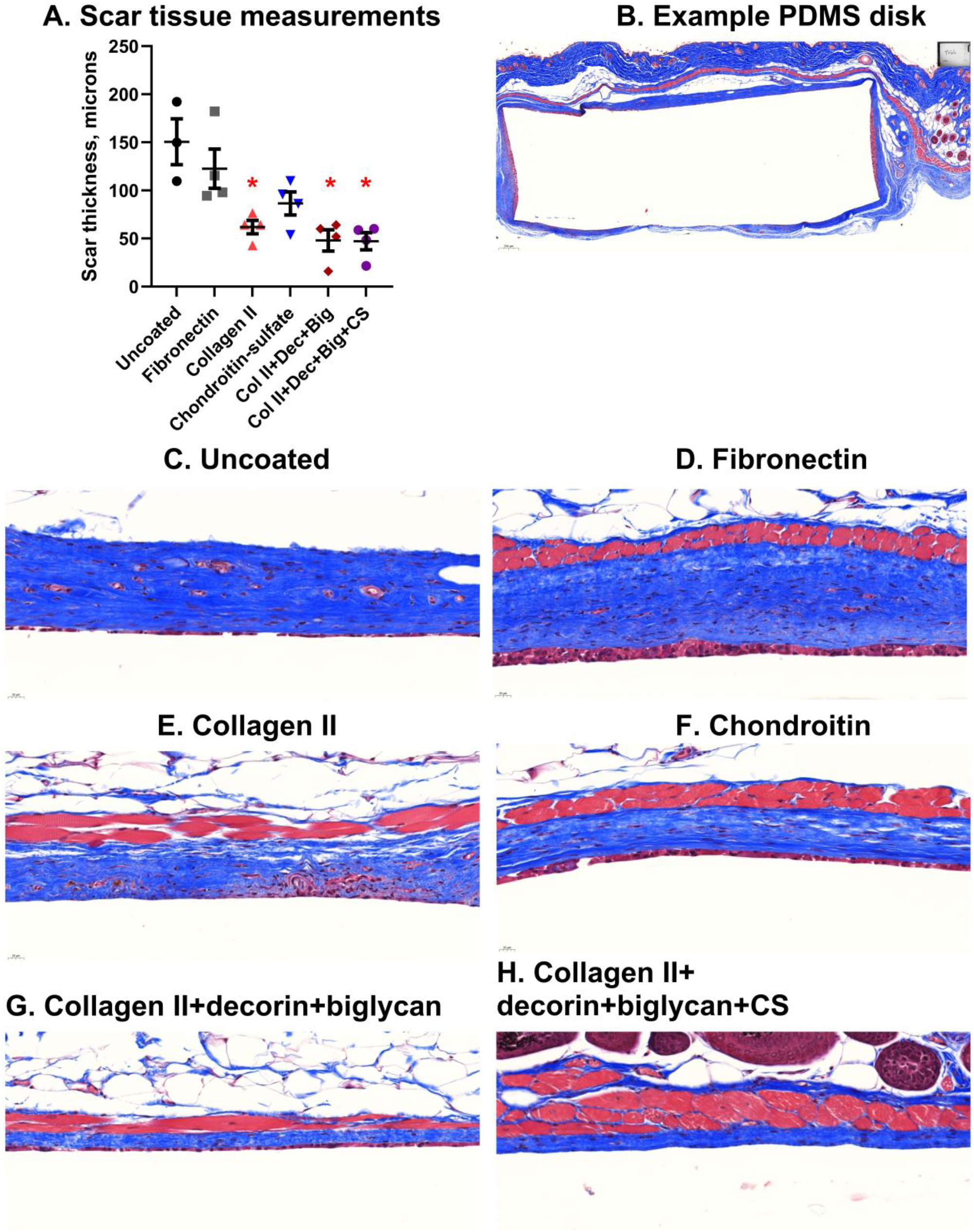
Coating of PDMS implants by the indicated ECM proteins reduces the scar tissue capsule thickness resulting from the foreign body response. 5 mm PDMS disks were sterilized and coated with 200 ug/ml of the indicated ECM proteins for 24 hours by adsorption. Coated PDMS disks were then implanted subcutaneously in the backs of C57Bl6 mice, and allowed to remain for 6 weeks. Resected skin (containing the implant) was paraffin-fixed and stained with Masson’s Trichrome to reveal scar tissue. (A) The thickness of scar tissue surrounding the resected implants was measured at the skin-implant interface (top of images B-H) at 5 different positions, and then averaged. n of at least 3, mean ± SEM * = statistical significance of P < 0.05, < 0.01, or < 0.001, significance vs uncoated PDMS disk control, 1-way ANOVA with Dunnett’s test for multiple comparisons. Panel A is significant by 1-way ANOVA with a post-test for trend between datasets read in a left-to-right direction. (B) A representative image of resected skin (top of the image) and the scar tissue (Blue) surrounding the position where the uncoated PDMS insert was located. PDMS dissolved during paraffin-fixation, but the dimensions of the PDMS disk are approximately retained by the fixed tissue, as seen in panel B. Size bar is 200 µm. (C-H) Representative images of the skin-PDMS interface, with scar tissue colored blue. Representative images are from areas of scar tissue without folding or separation of scar tissue layers. Size bar is 20 µm.

First, we coated a 3 mm polydimethylsiloxane (PDMS) disk by adsorption overnight under sterile conditions. We then implanted these disks subcutaneously into mice and resected the disks 6 weeks later, measuring the resulting scar tissue thickness (Fig 3A and 3B). Uncoated disks (Fig 3C) and disks coated with fibronectin (Fig 3D) were used as comparison controls for disks coated with collagen II (Fig 3E); chondroitin sulfate (Fig 3F); collagen II and decorin and biglycan (Fig 3G); and collagen II, decorin, biglycan, and chondroitin sulfate (Fig 3H). Disks were not coated with decorin or biglycan alone since these proteins together comprise only 15% of the mass of cartilage, while collagen II and chondroitin make up the majority of the remaining 85% of the mass. Therefore, disks coated with 100% biglycan or decorin would not be biologically relevant or help us elucidate why cartilage does not become fibrotic under healthy conditions. PDMS disks coated with fibronectin and chondroitin sulfate had an average surrounding scar thickness similar to the control PDMS disks. Disks coated with collagen II had significantly thinner surrounding scar tissue on average, as did disks coated with the mixture of collagen II, decorin, and biglycan (Fig 3A and 3G), as well as disks coated with the mixture of collagen II, decorin, biglycan, and chondroitin sulfate (Fig 3A and 3H).

In addition to adsorbing proteins to the surface of PDMS disks (Fig. 3), we wanted to attach proteins more stably to the surface of an implant via covalent bonding. For this experiment, we acquired 3 mm polyballs coated with a surface composed of carboxylate groups which covalently bind proteins in solution via EDAC-mediated chemistry. We coated these polyballs with the same amount and ratio of proteins (collagen II, decorin, biglycan, chondroitin sulfate) as we coated the PDMS disks in Fig. 3. We implanted these coated polyballs into mice subcutaneously and resected them 6 weeks later. Polyball implants generated less thick scar tissue compared to the PDMS implants (Fig. 4). As with the PDMS disks, comparison groups were uncoated polyballs, polyballs coated with fibronectin, and the same experimental coating groups (Fig. 4A-H). Compared to uncoated, coating with a mixture of collagen II, decorin, and biglycan, or a mixture of collagen II, decorin, biglycan, and chondroitin sulfate, caused thinner surrounding scar tissue (Figs 4G and H).

**Figure 4:**
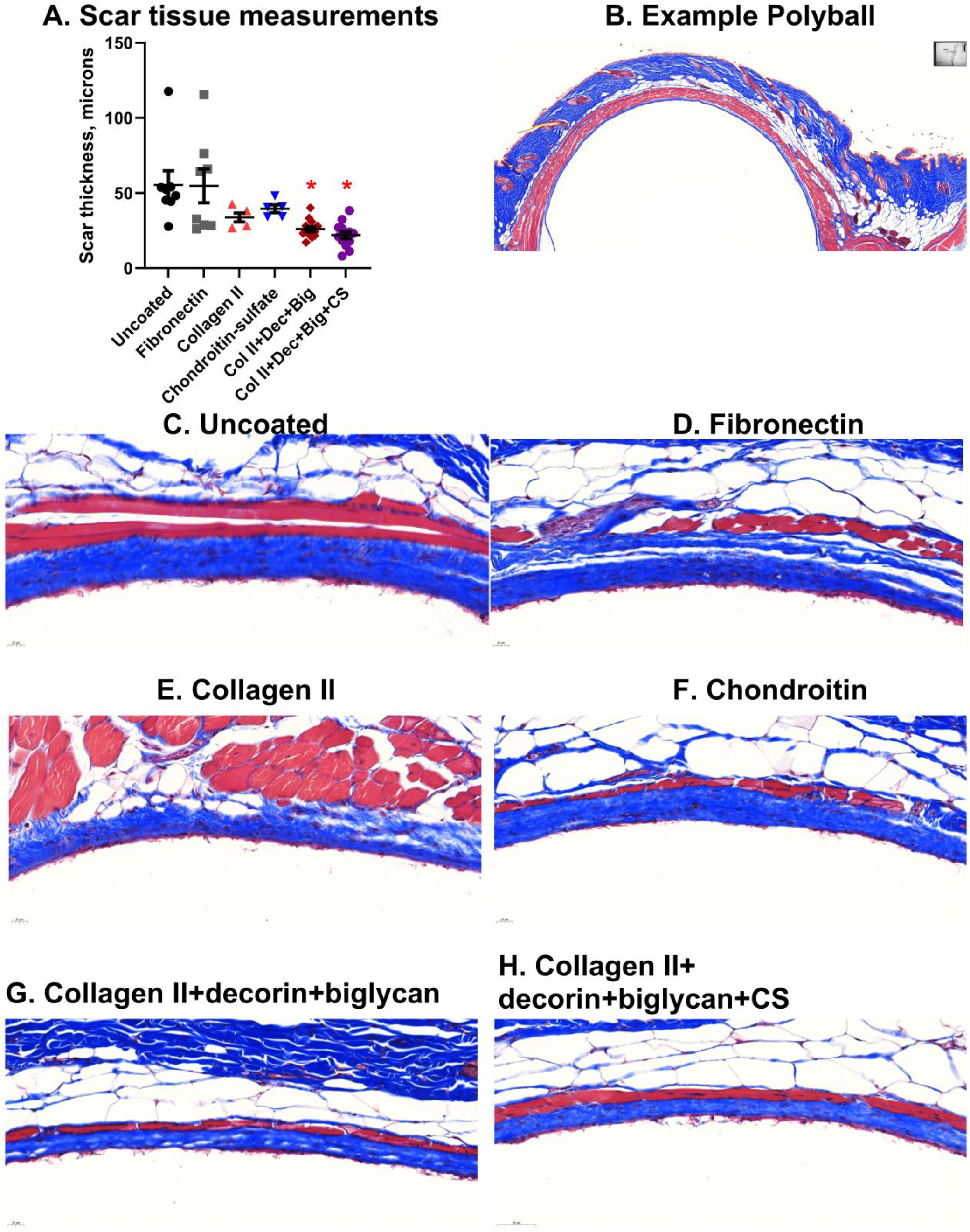
Coating of polyball implants by the indicated ECM proteins reduces the scar tissue capsule thickness resulting from the foreign body response. 3 mm polyballs were sterilized, and coated with 200 µg/ml of the indicated ECM proteins for 24 hours by covalent-coupling and adsorption. Coated polyballs were then implanted subcutaneously in the backs of C57Bl6 mice and allowed to remain for 6 weeks until sacrifice. Resected skin (containing the implant) was paraffin-fixed and stained with Masson’s Trichrome to reveal scar tissue. (A) The thickness of scar tissue surrounding the resected implants was measured at the skin-implant interface at 5 different positions, and then averaged (top of images B-H). Values are mean ± SEM, n=5 to 8. * P < 0.05 compared to uncoated PDMS disk control, 1-way ANOVA with Dunnett’s test for multiple comparisons Panel A is significant by 1-way ANOVA with a post-test for trend between datasets read in a left-to-right direction. (B) A representative image of resected skin (top of the image) and the scar tissue (Blue) surrounding the position where the polyball was located. Polyballs dissolved during paraffin-fixation, but the dimensions of the polyball disk are approximately retained by the fixed skin. Bar is 200 µm. (C-H) Representative images of the skin-polyball interface, with scar tissue colored blue. Size bar is 20 µm.

## Discussion

Our overall objective in this work is to explore the relationship between ECM environment and the initiation, progression, and maintenance of fibrotic scar tissue. We have previously investigated the role that mechanical stiffness in the cellular environment plays in the biology of fibrosis [48, 49]. In this report, we show that protein coating mixtures of collagen II, biglycan, decorin, and chondroitin sulfate (in proportions approximating the proportions found on the surface of articular cartilage) are anti-fibrotic, both *in vitro* and *in vivo*. Implants coated with this cartilage-like coating have less scar tissue deposition than uncoated implants or fibronectin-coated implants. We hypothesize that this reduction in scar-tissue deposition on the surface of implants may be due to cells’ ability to recognize complex ECM composition in the microenvironment.

Both the PDMS insert (Fig. 3) and the polyball (Fig. 4) remained intact during the formalin fixation of the resected tissue, but dissolved during the paraffin mounting of the tissue. This was advantageous for sectioning the tissues surrounding the implants because sectioning through PDMS or polystyrene was not necessary. However, dissolving the PDMS or polystyrene polyball left an open space within the scar tissue (visible in Fig. 3B and Fig. 4B). The lack of any support within the open space caused the layers of scar tissue to separate in places, or to fold (also visible in Fig. 3B and Fig. 4B). This necessitated averaging the scar tissue measurements from multiple locations around the implant that were unaffected by the tissue folding or the separation between scar tissue layers. The representative inset images in Figs. 3 (C-F) and 4 (C-F) are taken where there is little or no scar layers of scar tissue separation, or tissue folding. Muscle layer thickness was not consistent throughout the outer layer of the PDMS disk or polyball, and changes in the muscle layer thickness were not significantly different between treatments.

Several reports have analyzed the response ECM coatings have on myofibroblast differentiation, sometimes yielding conflicting results. Previous studies have found that collagen I appears to be anti-fibrotic when used as a coating [51]. Additionally, a peptide from the fibronectin “arginine-glycine-aspartate” (RGD) domain also inhibits myofibroblast differentiation when used as a coating [51]. Lumican [52], hyaluronic acid [53], and periostin [54] have also been shown to modulate myofibroblast differentiation. Conversely, the biology of how monocyte-derived myofibroblasts can bind and degrade ECM has also been explored [55].

Decorin has previously been shown to reverse scar tissue formation [56], and has also been shown to reduce (though not significantly) myofibroblast differentiation *in vitro* [52]. Chouhan *et al* [56] used a sustained release of decorin to reduce dermal scarring, indicating that the method of delivery may determine whether decorin reduces scar tissue formation when used as an anti-fibrotic. We deliver decorin as a pre-existing coated substrate, which differs from Chouhan. It is possible that decorin’s delivery method (at different time points or densities) may effect decorin’s interactions with collagen, and these differences could determine whether the added decorin is anti-fibrotic. For example, decorin may help in organizing newly initiated mass collagen deposition on a coated implant to decrease an overt fibrotic response. Alternatively, it may reinforce tense collagen bundles in an already fibrotic environment with mass collagen deposition, contributing to greater stiffness.

The finding that cartilage-like coatings resist fibrosis may have applications beyond the coating of implants. For instance, if the specific cellular receptors that engage cartilage are found, these receptors could then be targets for agonists or antagonists which may be effective anti-fibrotics. Cartilage is a stiff tissue with a Young’s modulus of 800 kPa [57] which suggests that the surface of cartilage would meet the minimum stiffness requirements of at least 12 kPa for scar tissue to form [48]. The results of our previous studies [48, 49] indicate that myofibroblast differentiation is dependent on at least two mechanical checkpoints. Myofibroblast precursors must be (1) adhered to a (2) surface of sufficient mechanical stiffness of at least 12 kPa. When myofibroblast precursors (monocytes of fibroblasts) are not adhered, or are adhered to a surface softer than 12 kPa, we were not able to potentiate the differentiation of myofibroblasts.

Because cartilage meets the minimum stiffness requirements for myofibroblast differentiation to occur, it raises the question of why cartilage is not a frequent substrate for the formation of scar tissue. We hypothesize that myofibroblast precursor cells assess not only whether their microenvironment meets minimum stiffness requirements, but also that myofibroblast precursor cells assess whether the ECM composition of their microenvironment is suitable for transition into myofibroblasts.

The results we present here show that myofibroblast differentiation is dependent not just on the mechanical environment, but also on the ECM microenvironment. Both microenvironmental stiffness and protein composition may act as checkpoints for the myofibroblast transition. Myofibroblast differentiation is governed by at least 3 mechanical/environmental checkpoints: Myofibroblast differentiation requires that myofibroblast precursors be (1) adhered (2) in an environment of suitable ECM composition and (3) sufficient mechanical stiffness of at least 12 kPa.

These findings could be used to develop more effective and stable coatings for implantable devices. Further studies could also address the use of different ECM compositions and stiffnesses used in engineered biomaterials. The clinical attractiveness of these findings depend on future studies exploring the interaction of cells with different ECM compositions of differing stiffnesses.

## Materials and Methods

### Study design

This study was designed to explore the relationship between the ECM environment and myofibroblast differentiation. Group size was selected based on experience with the pulmonary fibrosis model. Mice were randomized into treatment groups within a cage to eliminate cage effects from the experiment. Implants were also implanted and resected by multiple researchers to ensure reproducibility. All mouse experiments were performed under supervision with protocols approved by the University of Chicago IACUC (protocol 72450, approved 2/25/2020). Mice were housed in a 12 hour day/night cycle, with free access to food and water.

### Purification of primary human monocytes

All human blood was acquired through the University of Chicago blood donation center in accordance with a human subjects protocol at the University of Chicago (IRB15-0102). Human monocytes were purified as previously described [48]. Briefly, PBMCs were purified from leukocyte reduction filters from de-identified blood donors. Blood was filtered, and leukocytes purified, the same afternoon as a morning blood donation to reduce the amount of time that PBMCs could adhere to the filter.

The collected blood was layered with lymphocyte separation media (Corning, Corning, NY, 25-072-CI) and centrifuged at 1300 xg for 20 min. The PBMC layer was then removed by pipetting. This protocol regularly resulted in PBMC recovery of approximately 300 million cells per filter.

Monocytes were purified from PBMCs by use of a negative selection kit for human monocytes (Stemcell, Cambridge, MA), per the manufacturer’s instructions. Approximately 20 million monocytes were purified from each filter. Monocytes were then washed by PBS using five successive 300 x g centrifugation steps, to remove EDTA from the resulting population. Monocytes were checked for purity using flow cytometry via size assessment and forward and side scatter characteristics, and average purity was above 95%.

### Purification of primary mouse monocytes

All mouse experiments were performed under supervision with protocols approved by the University of Chicago IACUC. Mouse monocytes were purified as previously described [48]. Spleens were resected from healthy C57BL/6 mice, pooled, and were placed in PBS with 1 mM EDTA, 2% fetal calf serum. All PBS, plasticware, filters, glassware and magnets were pre-chilled to 4°C, and kept cold throughout this procedure to limit the clumping of cells.

Spleens were pushed through a 100 μm filter to disassociate the cells. Monocytes were purified from disassociated cells by use of a negative selection kit (Stemcell 19861), following the manufacturer’s instructions. The purified monocytes were then washed by PBS using five successive 300 x g centrifugation steps in order to remove EDTA from the resulting population. Monocytes were checked for purity using flow cytometry via size assessment and forward and side scatter characteristics, and average purity was above 95%. The average yield was 1.5 million monocytes per spleen.

### Culture of human and mouse monocytes

Human and mouse monocytes were cultured as previously described, using serum-free media (SFM) [48]. Briefly, SFM for human cells is composed of Fibrolife (Lifeline, Frederick, MD LM-0001), with 1x ITS-3 (Sigma, St. Louis, MO I2771), 1x HEPES buffer (Sigma), 1x non-essential amino acids (Sigma), 1x sodium pyruvate (Sigma), and penicillin-streptomycin with glutamate (Sigma). For mouse monocytes, 2x concentrations of ITS-3, HEPES buffer, non-essential amino acids, and sodium pyruvate were added, with 50 μM β-mercaptoenthanol (ThermoFisher, Waltham, MA, 21985023) and pro-fibrotic supplements M-CSF (25 ng/ml, Peprotech, Rocky Hill, NJ) and IL-13 (50 ng/ml) to induce myofibroblast differentiation. Additionally, M-CSF and IL-13 were refreshed in the media of mouse monocytes after 3 days of culture. Monocytes were allowed to differentiate for 5 days, and counted based on morphology as previously described [58].

Human monocytes were cultured immediately following purification, at 100,000 monocytes/cm^2^, and monocytes were cultured immediately following purification, at 250,000 monocytes/cm^2^.

### Culture of human and mouse fibroblasts

Human fibroblasts (MRC-5, ATCC, Manassas, VA) and mouse fibroblasts (NIH-3T3, ATCC) were cultured in SFM composed for human cells, with 1x concentrations of additives. 5 ng/ml TGFβ (Peprotech, Cranbury, NJ, 100-21) was added to induce myofibroblast formation [59]. Cells were cultured at 10,000/cm^2^, and TGFβ was refreshed in cultures weekly to maintain the myofibroblast phenotype, if necessary.

### Coating for ECM screen

Tissue-culture-treated plastic was coated with 10 ug/ml of human ECM proteins for 1 hr at 37 degrees or overnight at 4 degrees and washed twice with 1X PBS. Cells were plated onto fresh ECM that had never dried, similar to previous ECM-coating experiments [60]. ECM components were acquired from various companies. All collagens were from Millipore (C4243, C7774, CC052, CC054, CC076), all laminins were from BioLamina (LN111, LN121, LN211, LN221, LN332, LN411, LN421, LN511), fibronectin (Sigma-Aldrich F0895), vitronectin (Sigma-Aldrich, CC080), tenascin C (Millipore CC065), osteopontin (Sigma-Aldrich, SRP3131), fibrinogen (Enzyme Research Laboratories, FIB 3), chondroitin sulfate (Sigma-Aldrich C9819), decorin and biglycan were from SinoBio (10189-H08H and 10447-H08H).

### Coating of PDMS inserts

PDMS inserts were sterilized by exposure to UV light, and were coated by adsorption in a PBS solution with 200 ug/ml of overall protein (100 ug/ml collagen II, 70 ug/ml chondroitin sulfate, 15 ug/ml decorin and 15 ug/ml biglycan, matching relative concentrations of collagen II (50%), decorin (7.5%), biglycan (7.5%), and chondroitin sulfate (35%) in the superficial level of articular cartilage [35]. PDMS inserts were coated overnight and were dipped thrice in PBS before insertion into mice to remove un-adsorbed proteins.

### Coating of Polyballs

Polyballs (Polysciences 19841) were coated according to the manufacturer’s instructions, using a Polylink protein coupling kit (Polysciences 24350). Briefly, polyballs coated with carboxyl groups (-COOH) were exposed to 1-ethyl-3-(3-dimethylaminopropyl) carbodiimide hydrochloride (EDAC), which creates an active ester that is reactive towards primary amines. We then placed these polyballs into 200 ug/ml of overall protein (100 ug/ml collagen II, 70 ug/ml chondroitin sulfate, 15 ug/ml decorin and 15 ug/ml biglycan, matching relative concentrations of collagen II (50%), decorin (7.5%), biglycan (7.5%), and chondroitin sulfate (35%) in the superficial level of articular cartilage [35]. Polyballs were coated overnight and were dipped thrice in PBS before insertion into mice to remove un-adsorbed proteins.

### Flow cytometry

Cultured myofibroblasts were lifted with ice cold trypsin-EDTA (Sigma T4049) followed by mechanical agitation by a rubber policeman. Myofibroblasts were fixed and permeabilized using using Cytofix/Cytoperm (BD biosciences, Franklin Lakes, NJ 554722), and live-dead stained (live-dead aqua, ThermoFisher, L34957) per manufacturer’s instructions. Antibodies used were α-collagen I (Biolegend, San Diego, CA, FITC), anti-α smooth muscle actin (αSMA) (R and D systems, Minneapolis, MN BV421), α-ki-67 (BD biosciences, PE). Compensation was performed via UltraComp beads (ThermoFisher) per the manufacturer’s instructions.

### Legendplex cytokine detection by ELISA

Frozen supernatant from cultured human and mouse myofibroblasts were thawed at 4°C and centrifuged at 4°C and 10,000 x g to pellet cell debris. Supernatant was taken and added to 96 well round bottom plates, in duplicate. Legendplex (Biolegend 740809) beads against general inflammation markers were added, according to the manufacturer’s instructions. For human monocytes, sample readouts were normalized to each individual donor control. Donor controls were not used for pooled mouse monocytes or immortal fibroblasts.

### Implant insertion

Mice were anesthetized via isoflurane inhalation (2%) and subcutaneously injected with meloxicam (1 mg/kg), buprenorphine (0.1 mg/kg) in a saline solution. Mice were shaved, and a small incision made between the shoulder blades. 4 Polyballs were inserted through this incision to different locations on the mouse’s back, using a long trocaring needle. PDMS implants were inserted through this incision using thin sterile forceps. The incision was closed using wound clamps and mice were checked daily for signs of distress.

### Implant resection

Implants were resected from mice 6 weeks after implantation. Mice were euthanized by CO_2_ inhalation. Implants were located by palpitating the skin. After locating the implants, the whole section of skin was removed and placed in a histology cassette with 2 sponges above and below and then placed directly into 10% neutral buffered formalin.

### Histology

Implant resections were placed in 10% neutral buffered formalin for 5 days 4°C and then mounted in paraffin. The process of paraffin mounting dissolved the styrene-based polyballs and the PDMS inserts, which is why no implants are visible in the histology images (Figs. 3B and 4B).

Implants were sectioned into 5 mm slices and stained with Masson’s trichrome via a Bond-Max autostaining system (Leica biosystems, Lincolnshire, IL) in order to visualize the collagen-rich scar tissue.

### Microscopy and analysis of scar tissue thickness

Stained implant resections were scanned at high resolution using a CRi Panoramic SCAN 40x Whole Slide Scanner (Perkin-Elmer). Thickness was measured using ImageJ, and at least 5 thicknesses were measured and averaged to ensure that the thickness of scar tissue was adequately measured.

## Statistical analysis

Statistical analyses were performed using GraphPad Prism software, and *P* < 0.05 was considered statistically significant. Two different statistical analyses were used: 1-way ANOVA with Dunnett’s post test for multiple comparisons, or 1-way ANOVA test for test for linear trend between dataset mean and left-to-right order (which was used to compare the trend relationships in Figs. 1, 2, 3 and 4, and supplemental Figs. 2 and 3).

## Supporting information

Figure 4 at highest possible resolution for BioRXiv upload limit

## Acknowledgements

We thank the Human Tissue Resource Center of the University of Chicago for histology analysis. We thank the Integrated Light Microscopy Core of the University of Chicago for Imaging. We’d also like to acknowledge the staff of the animal resources center at the University of Chicago, particularly Ani Solanki, who invented several minimally invasive insertion methods of these implants using only a single incision.

## Abbreviations

(αSMA): alpha-smooth muscle actin
(RGD): arginine-glycine-aspartate domain
(CCL22): C-C Motif Chemokine Ligand 22
(CCL17): C-C Motif Chemokine Ligand 17
(CXCL1): chemokine ligand 1
(ELISA): enzyme-linked immunoassay
(ECM): extracellular matrix
(FPLC): fast protein liquid chromatography
(GAG): glycosaminoglycan
(MRC-5) Interleukin 6 (IL6): Human fibroblasts
(IL-12p70): Interleukin 12 subunit p70
(IL13): Interleukin 13
(M*-*CSF): Macrophage colony-stimulating factor
(NIH-3T3): mouse fibroblasts
(PBMC): peripheral blood mononuclear cells
(PBS): phosphate buffered saline
(PDMS) serum-free media (SFM): Polydimethylsiloxane
(SLRP): small leucine-rich proteoglycan
(TGFβ): Transforming growth factor β
(TLR2): toll-like receptors 2
(TLR4): toll-like receptors 4
(TNFα): Tumor necrosis factor α
(EDAC): 1-ethyl-3-(3-dimethylaminopropyl)carbodiimide hydrochloride

## Funding

This work was supported in part by the University of Chicago (to JAH) and the rebuilding the kidney consortium (RBK, to JAH)

## Author’s contributions

Conceptualization: MJVW, MO, RHG, DP

Methodology: MJVW, MO, RHG, DP

Investigation: MJVW, MO, AS, DP

Visualization: MJVW

Funding acquisition: JAH, RHG

Project administration: MJVW, JAH

Supervision: MJVW, JAH

Writing – original draft: MJVW

Writing – review & editing: MO, JAH, RHG, DP

## Ethics approval

All the animal experiments performed in this work were approved by the Institutional Animal Care and Use Committee of the University of Chicago.

## Conflicts of interest

The authors declare that they have no competing interests.

## Notes

### Competing Interest Statement

The authors have declared no competing interest.

