## Supplementary Figures for "An extracellular matrix coating similar to articular cartilage inhibits the differentiation of myofibroblasts and limits the development of implant-induced fibrosis"

**Supplemental Figures**

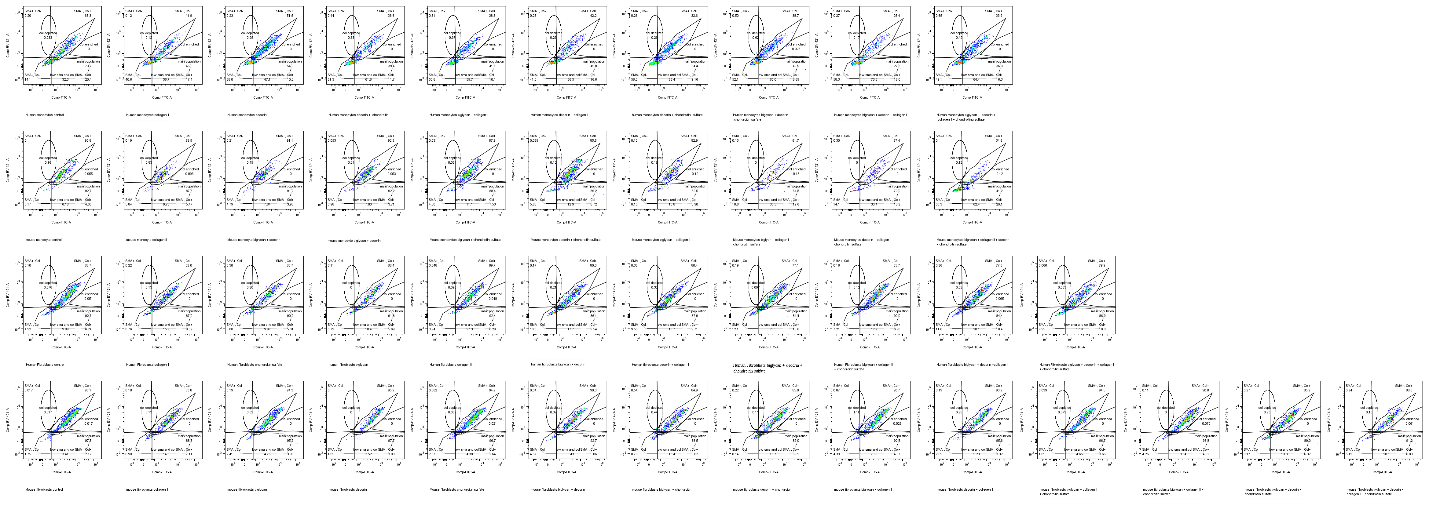

Below D) mouse fibroblasts

C) Human fibroblasts

B) Mouse monocytes

A) Human monocytes

**Supplemental Figure 1: Representative flow plots from a single experiment showing the percentage of the population that αSMA and collagen I double positive.** Values were normalized to controls and can be found in Fig. 2.

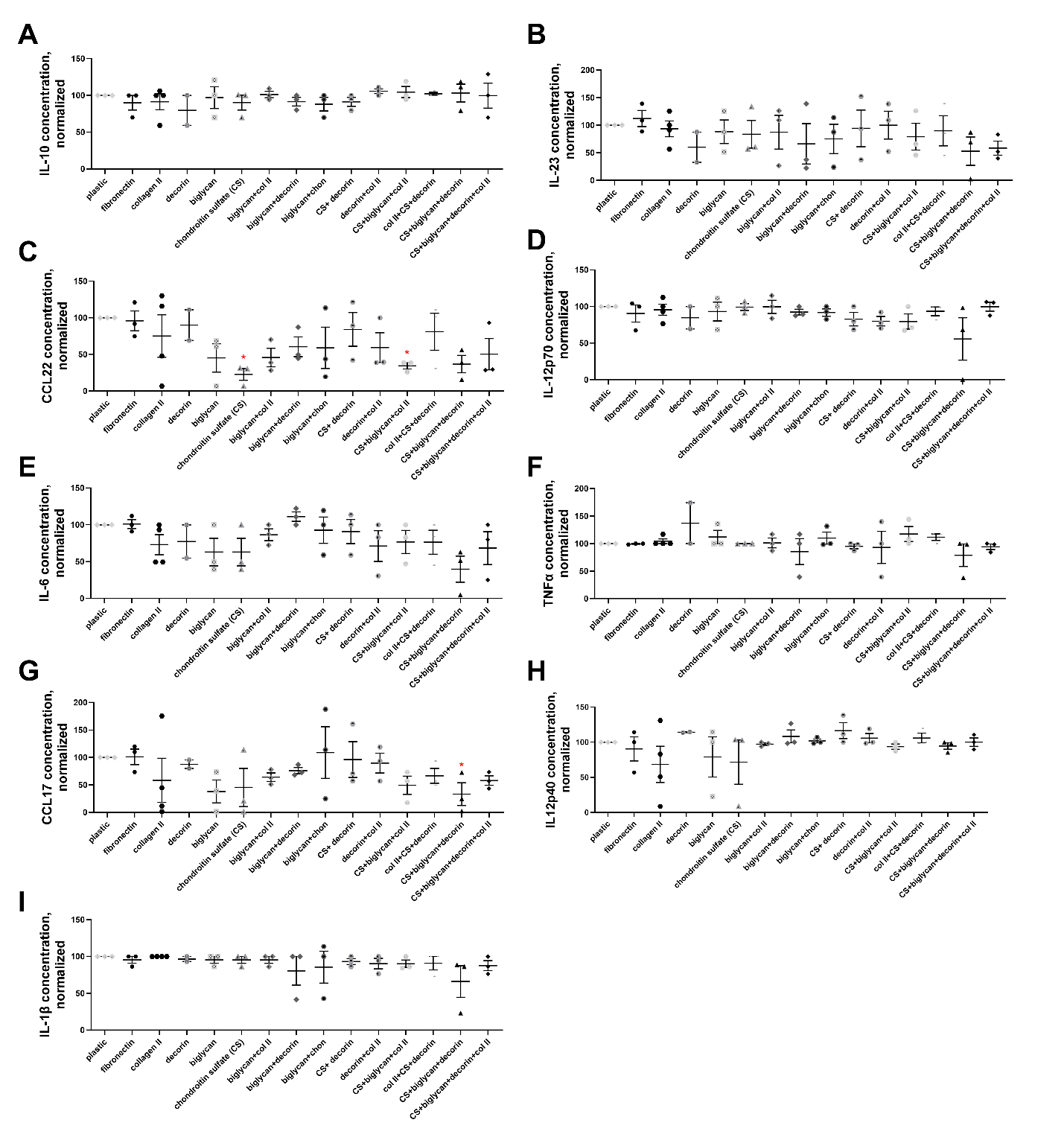

**Supplemental Figure 2: Culture on ECM proteins comprising the main protein components of cartilage reduces mouse monocyte’s pro-fibrotic secretome.** Freshly isolated mouse monocytes were cultured for 5 days under pro-fibrotic conditions (M-CSF and Il-13) on surfaces coated with ECM proteins. Conditioned media from the culture was assessed via Legendplex ELISA. Only results over the detection limit are displayed here. Secreted proteins over the detection limit were: (A) IL-10, (B) IL-23, (C) CCL22, (D) IL-12 subunit p70, (E) IL-6, (F) TNF-α, (G) CCL17, (H) IL-12 subunit p40, and (I) IL-1β. n = 3. * = statistical significance of P < 0.05, < 0.01, or < 0.001, significance vs uncoated tissue-culture surface control, 1-way ANOVA with Dunnett’s test for multiple comparisons. 1-way ANOVA test for trend for datasets means from left to right is significant for C (CCL22, **), D (IL12-p70, *), E (IL-6 ,*), with no significance for trend for any other panel.

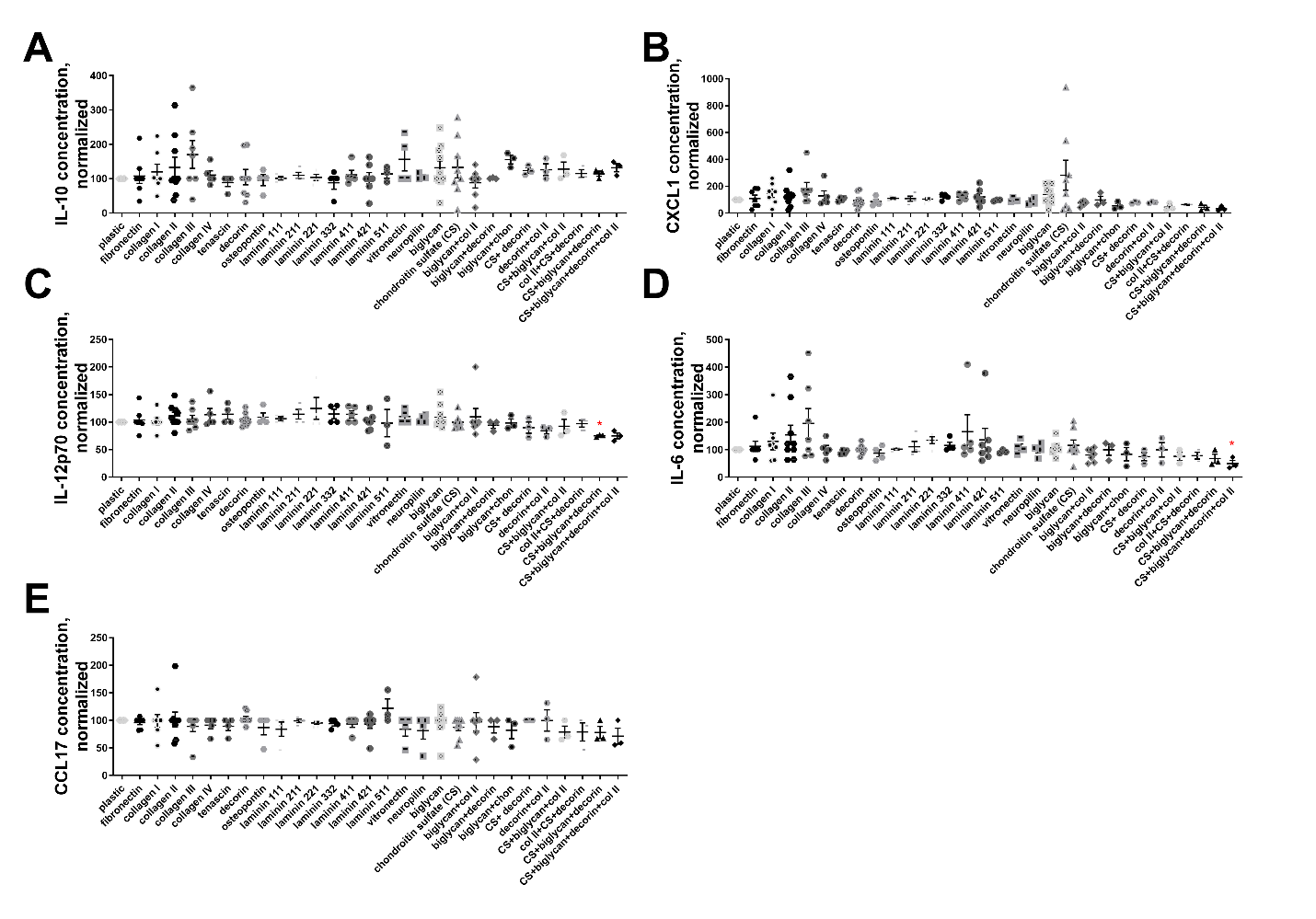

**Supplemental Figure 3: Culture on ECM proteins comprising the main protein components of cartilage reduces mouse fibroblast’s pro-fibrotic secretome.** Mouse fetal fibroblasts were cultured for 5 days under pro-fibrotic conditions (TGFβ) on surfaces coated with ECM proteins. Conditioned media from the culture was assessed via Legendplex ELISA. Only results over the detection limit are displayed here. Secreted proteins over the detection limit were: (A) IL-10, (B) CXCL1, (C) IL-12 subunit p70, (D) IL-6, and (E) CCL17. n ranges from 3 to 5 . * = statistical significance of P < 0.05, < 0.01, or < 0.001, significance vs uncoated tissue-culture surface control, 1-way ANOVA with Dunnett’s test for multiple comparisons. 1-way ANOVA test for trend for datasets means from left to right is not significant for A (IL-10), but is significant for B (CXCL1, *), C (IL12-p70, ***), D (IL-6, **), and E (CCL17, *).
