## Supplementary figures and images for "An extracellular matrix coating similar to articular cartilage inhibits the differentiation of myofibroblasts and limits the development of implant-induced fibrosis"

### Figure 4 at highest possible resolution for BioRXiv upload limit

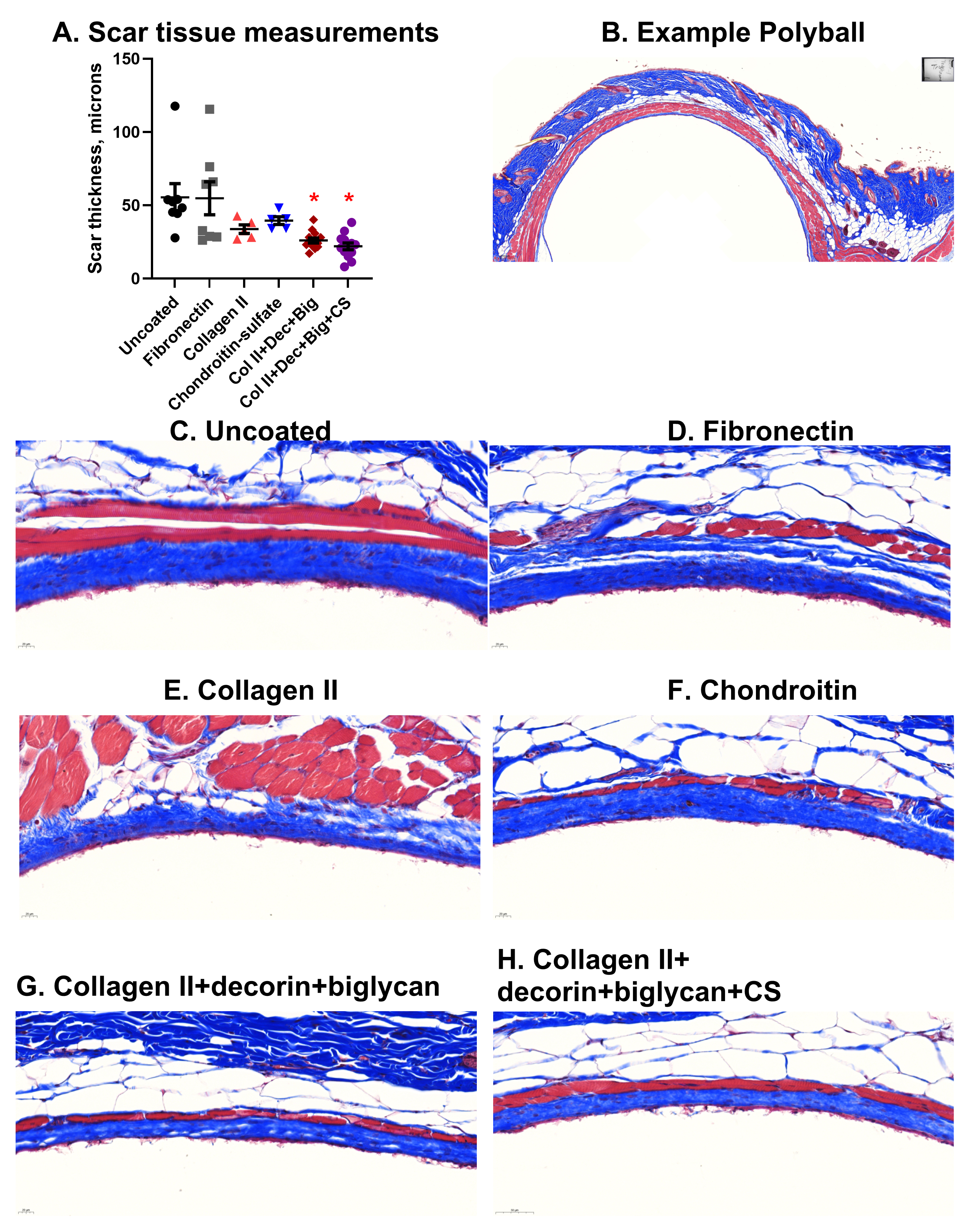
